# DeepChIA-PET: Accurately predicting ChIA-PET from Hi-C and ChIP-seq with deep dilated networks

**DOI:** 10.1101/2022.10.19.512935

**Authors:** Tong Liu, Zheng Wang

**Affiliations:** Department of Computer Science, University of Miami, 1365 Memorial Drive, P.O. Box 248154, Coral Gables, FL, 33124, USA

## Abstract

Chromatin interaction analysis by paired-end tag sequencing (ChIA-PET) can capture genome-wide chromatin interactions mediated by a specific DNA-associated protein. The ChIA-PET experiments have been applied to explore the key roles of different protein factors in chromatin folding and transcription regulation. However, compared with widely available Hi-C and ChIP-seq data, there are not many ChIA-PET datasets available in the literature. A computational method for accurately predicting ChIA-PET interactions from Hi-C and ChIP-seq data is needed that can save the efforts of performing wet-lab experiments. Here we present DeepChIA-PET, a supervised deep learning approach that can accurately predict ChIA-PET interactions by learning the latent relationships between ChIA-PET and two widely used data types: Hi-C and ChIP-seq. We trained our deep models with CTCF-mediated ChIA-PET of GM12878 as ground truth, and the deep network contains 40 dilated residual convolutional blocks. We first showed that DeepChIA-PET with only Hi-C as input significantly outperforms Peakachu, another computational method for predicting ChIA-PET from Hi-C but using random forests. We next proved that adding ChIP-seq as one extra input does improve the classification performance of DeepChIA-PET, but Hi-C plays a more prominent role in DeepChIA-PET than ChIP-seq. Our evaluation results indicate that our learned models can accurately predict not only CTCF-mediated ChIA-ET in GM12878 and HeLa but also non-CTCF ChIA-PET interactions, including RNA polymerase II (RNAPII) ChIA-PET of GM12878, RAD21 ChIA-PET of GM12878, and RAD21 ChIA-PET of K562. In total, DeepChIA-PET is an accurate tool for predicting the ChIA-PET interactions mediated by various chromatin-associated proteins from different cell types. DeepChIA-PET is publicly available at http://dna.cs.miami.edu/DeepChIA-PET/.

## 1 Introduction

Chromatin interaction analysis by paired-end tag sequencing (ChIA-PET) (Fullwood, et al., 2009) is a technique that processes chromatin immunoprecipitation (ChIP)-enriched chromatin complexes by linker ligation, proximity ligation, and high-throughput sequencing to identify significant long-range chromatin interactions at the whole genome. Compared with Hi-C (Lieberman-Aiden, et al., 2009), the main advantage of ChIA-PET is that the interactions it captured are located at the binding sites of one specific DNA-associated protein of interest, such as the insulator binding protein CTCF (Tang, et al., 2015), the RAD21 subunit of the cohesin complex (Grubert, et al., 2020; Heidari, et al., 2014), and RNA polymerase II (RNAPII) (Li, et al., 2012).

The CTCF-mediated and RAD21-mediated ChIA-PET experiments can be applied to investigate their key roles in chromatin folding and the establishment of topologically associating domains (Dixon, et al., 2012). The RNAPII-mediated ChIA-PET interactions are an excellent source of studying transcription regulation. The genome-wide chromatin interactions captured by the Hi-C technique and its variants (Rao, et al., 2014) can be thought of as a pool of ChIA-PET interactions including all chromatin-associated proteins, from which we can hardly identify interactions that are only related to one specific protein factor. Therefore, ChIA-PET is more applicable than Hi-C if exploring the potential functions of different DNA-associated proteins in the three-dimensional (3D) genome is of interest.

The chromatin immunoprecipitation followed by high-throughput sequencing (ChIP-seq) is a widely used method for analyzing interactions between DNA and chromatin-associated proteins, which can provide the binding sites of protein factors. Technically, we can think of ChIA-PET as a hybrid of Hi-C and ChIP-seq. However, there are not enough experimental ChIA-PET datasets publicly available in the literature; only 10 ChIA-PET experimental sets are available on the 4D Nucleome website (Dekker, et al., 2017) compared with 335 Hi-C and 141 ChIP-seq sets (as of 10/07/2022). Therefore, computationally predicting ChIA-PET interactions from Hi-C and ChIP-seq is a promising way to enrich ChIA-PET datasets.

There are some computational methods in the literature designed for processing ChIA-PET data and directly detecting peaks, such as Mango (Phanstiel, et al., 2015), cLoops (Cao, et al., 2020), and ChIAMM (Arega, et al., 2020). There are also some computational methods for predicting CTCF-mediated chromatin loops from DNA sequence-based features, such as CLNN-loop (Zhang, et al., 2022) and CTCF-MP (Zhang, et al., 2018), from various genomic and epigenomic features, such as Lollipop (Kai, et al., 2018), and from CTCF and Hi-C data, such as LOOPbit (Galan, et al., 2022). However, none of these computational methods are targeted for learning and predicting ChIA-PET interactions.

Loop-Extrusion-Model (Xi and Beer, 2021) used a simple mathematical model of CTCF-mediated loop formation for predicting CTCF ChIA-PET loops. The limitations of this method are that the authors did not blindly test on non-CTCF ChIA-PET and widely available Hi-C data are not used as reference input. Peakachu (Salameh, et al., 2020) overcame the second limitation by directly using Hi-C to predict ChIA-PET interactions. However, Peakachu has the following disadvantages: (1) only Hi-C data as input may result in that the learned models are ChIP-specific. This is because bulk Hi-C data is not protein specific. Therefore, the machine learning model is trained to map non-protein-specific Hi-C data to protein-specific ChIA-PET interactions. Next time when the same machine learning model is used to predict the ChIA-PET interactions formed by a different type of protein, the input will be the same, which may cause the problem that the machine learning model may only be adapted to or work well for the type of ChIA-PET data used for training. However, adding the ChIP-seq data in input as we have done in this research solves this problem to some extent. (2) the receptive field of machine learning was restricted to a 11 × 11window for representing the Hi-C features of the center pixel; (3) the number of negative samples for training was set equal to the number of positive pixels, resulting in that millions of unselected negative pixels in two-dimensional (2D) Hi-C contact matrices are never seen by the machine learning method (i.e., random forest); and (4) the strategy of one positive pixel as an independent training sample sacrifices the valuable information that one anchor may involve in multiple long-range ChIA-PET interactions.

In this paper, we present DeepChIA-PET, a deep-learning method for accurately predicting ChIA-PET interactions based on Hi-C and ChIP-seq data. DeepChIA-PET applied a deep dilated, residual convolutional network to learn the mapping from Hi-C and ChIP-seq to ChIA-PET at 10-kb resolution, which makes receptive fields larger enough for capturing long-range interactions and lets the deep learning method see all positive and negative pixels. Our evaluation results indicate that DeepChIA-PET significantly outperforms Peakachu, and our models trained with CTCF ChIA-PET can be accurately applied to predict non-CTCF ChIA-PET.

## 2 Materials and methods

### Data processing for Hi-C, ChIP-seq, and ChIA-PET

We downloaded KR-normalized Hi-C data for three different cell types (GM12878, HeLa, and K562) at 10-kb resolution using Juicer (Durand, et al., 2016) (Table S1), which are also available on Gene Expression Omnibus (GEO) under accession number GSE63525 (Rao, et al., 2014). Hi-C peaks in human GM12878 were detected with HiCCUPS (Rao, et al., 2014) at 10-kb resolution. TAD annotations of GM12878 at 10-kb resolution were downloaded from TADKB (Liu, et al., 2019), which used the directionality index (DI) (Dixon, et al., 2012) to call TADs.

The ChIP-seq data for three different DNA-associated proteins (CTCF, RNAPII, and RAD21) were downloaded from UCSC website (Table S2). The average value for each 10-kb bin was calculated by pyBigWig (https://github.com/deeptools/pyBigWig).

We used various ChIA-PET datasets in this study (Table S3). The CTCF-mediated and RNAPII-mediated ChIA-PET interactions in GM12878 and HeLa were downloaded from GEO under accession number GSE72816. The RAD21-mediated ChIA-PET interactions for GM12878 and K562 were obtained from Table S1 in (Heidari, et al., 2014). Since the anchor regions of ChIA-PET interactions are not binned at a given resolution, we assign one (positive) to pixels in a 2D ChIA-PET contact matrix at 10-kb resolution if the pixels overlap with two anchor regions of any known ChIA-PET interactions, and zero otherwise. Since the lengths of almost all anchors are less than 10,000 bp (Fig. S1A), the number of positive pixels is usually somewhat fewer than the number of raw ChIA-PET interactions.

The resolution is fixed to 10 kb for all three data types (i.e., Hi-C, ChIP-seq, and ChIA-PET) in this study. The Hi-C samples for training, validation, and blind test were extracted along the diagonal of a 2D contact matrix with a sliding window of 250 × 250 and a step of 50 bins. The ChIP-seq samples were obtained along each chromosome with a sliding window of 250 bins and a step of 50 bins. Since the genomic distances of almost all ChIA-PET interactions are less than 2 Mb (Fig. S2B), the genomic distance we covered by 250 bins (≤ 2.5 Mb) is long enough.

For each Hi-C contact matrix, we first rescaled the contacts by log transformation of log_2_(*x* + 1) and then further rescaled all contacts to the range [0,1] by min-max normalization. We calculated the mean and standard deviation (SD) of all Hi-C samples to perform final z-score normalization for each rescaled Hi-C matrix. For normalizing ChIP-seq vectors, we first did min-max normalization and then did z-score normalization as in normalizing Hi-C. The means and SDs for normalizing Hi-C and ChIP-seq were obtained from Hi-C and CTCF ChIP-seq of GM12878 and used on all the other testing datasets.

### Training, validation, and blind test

Our deep models were trained with CTCF-mediated ChIA-PET of GM12878 as ground truth. For blindly testing on each chromosome from 1 up to X, we trained 23 separate models. Specifically, excluding the current testing chromosome, the validation data were extracted from the largest chromosome and the rest chromosomes were used for generating the training data. The 23 learned models were also used for predicting non-CTCF ChIA-PET of a different cell line.

### DeepChIA-PET architecture

We illustrated the pipeline of DeepChIA-PET in Fig. 1. The original inputs are 2D Hi-C contact matrices and one-dimensional (1D) ChIP-seq vectors. To feed 1D ChIP-seq into our 2D convolutional network, we first converted the 1D ChIP-seq data into 2D by copying the ChIP-seq vector column- and row-wise, resulting in two 2D ChIP-seq matrices. We then concatenated the Hi-C and ChIP-seq matrices and obtained a 3D tensor (250 × 250 × 3) as input.

**Fig. 1.**
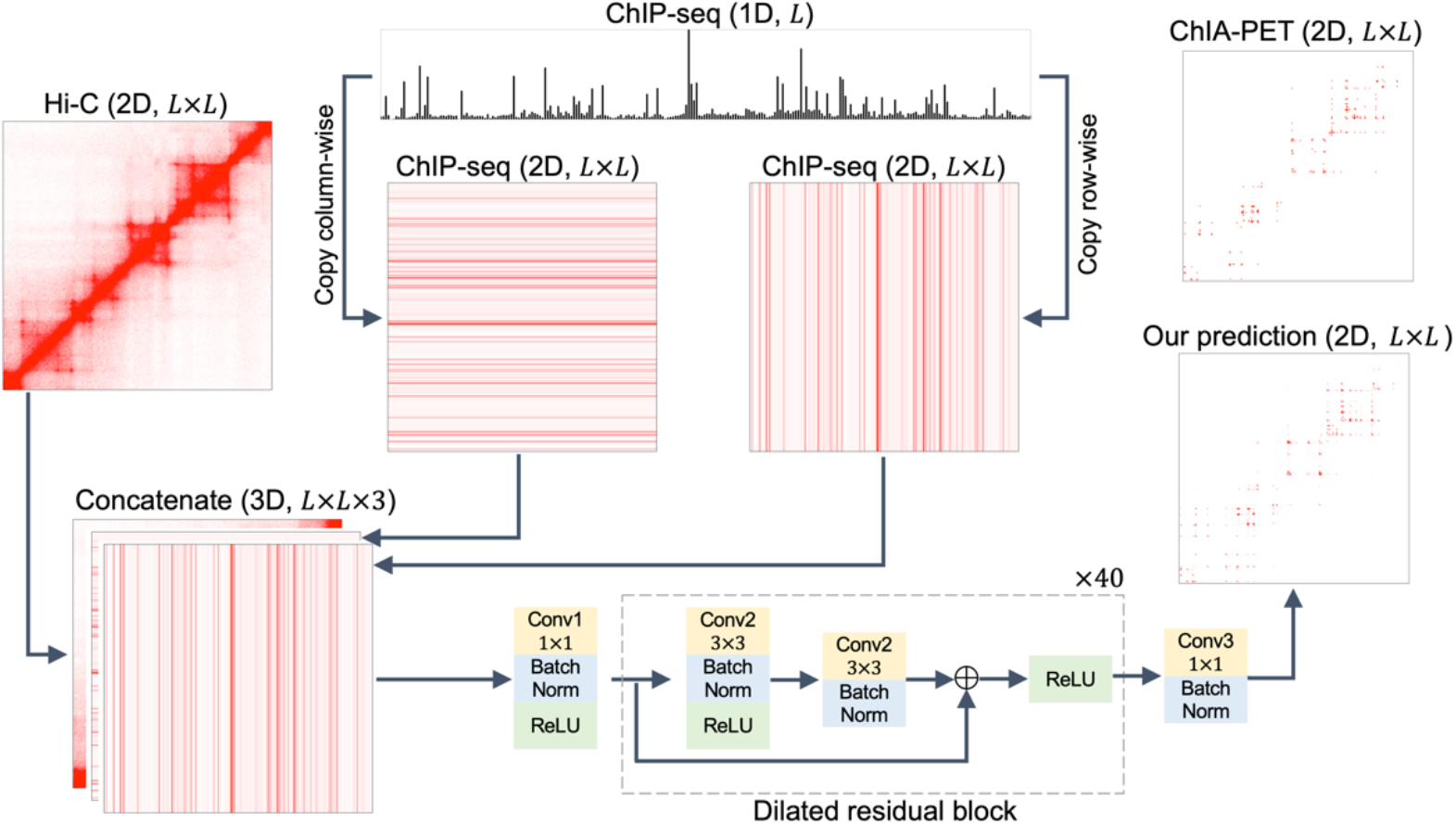
Framework of DeepChIA-PET. The 1D sequential ChIP-seq data are converted into two 2D pairwise matrices, which are further concatenated with Hi-C. The dashed box contains a typical residual block.

The final deep network we used is inspired by two deep networks for the prediction of protein contact maps (Kandathil, et al., 2019; Wang, et al., 2017) and contains three main parts. The first part consists of a 2D convolutional layer (Conv1 with 1 × 1 kernel size) for enhancing the hidden dimension from three to 128, a batch normalization layer, and a ReLU (Nair and Hinton, 2010). The second part consists of 40 typical residual blocks (He, et al., 2016) for learning the latent relationship between ChIA-PET and our inputs. Each block contains two dilated 2D convolutional layers (Conv2 with 3 × 3 kernel size) (Yu and Koltun, 2015). We set different dilation values for the two Conv2 layers in each block for capturing multi-scale, long-range ChIA-PET interactions. The last part consists of a 2D convolutional layer (Conv3 with 1 × 1 kernel size) and a batch normalization layer for reducing the output channel from 128 to one.

Since we may predict one pixel more than one time, the predicted score for the pixel is the average value of all its predicted scores. Moreover, since ChIA-PET interaction matrices should be symmetric, we further averaged the predicted values in the upper and lower triangular in matrices to get the final predicted scores. After obtaining the final predicted score for each pixel, we sort all predicted pixels by their predicted probabilities in descending order.

### Implementation details

We implemented our network in PyTorch (Paszke, et al., 2019). The optimizer we used was Adam (Kingma and Ba, 2014) with a weight decay of 1e-04. We tested five different batch sizes (4, 8, 16, 32, and 64) and used two strategies for setting the learning rate. The first one is to use a fixed value (0.01, 0.001, and 0.0001), and the second one is to dynamically adjust the learning rate by initially setting it to 0.001 and then reducing it with a factor of 0.1 when the validation loss stops improving for ten epochs. We tested two kernel sizes (3 and 5), two normalization methods (batch and instance), and two hidden dimensions (64 and 128). The number of residual blocks and the corresponding dilations for each block that we tested can be found in Table S4. The loss function for all models is binary cross entropy. We also used different positive weights (1, 3, 6, 9, and 12) when calculating loss. The best model for the blind test is the model that achieved the lowest validation loss. All models were trained in parallel on four NVIDIA A100 GPUs; each is equipped with 40 GB of memory.

### Evaluation metrics

Because the genomic distances of ground-truth ChIA-PET interactions are within a certain range, we only evaluated the pixels that are within the genomic range from 20 kb to 2 Mb. Since the ratios between the numbers of negative and positive pixels are usually larger than 500 (Fig. S2), we defined five different negative pixel sets (neg=pos, neg=5pos, neg=20pos, and neg=100pos, and neg=All) for evaluation. The number of negative pixels in the first four sets for each chromosome was set to the given number times the number of positive pixels on this chromosome, and the negative pixels for evaluation were randomly selected from all negative pixels. In the last negative set (neg=All), we used all negative pixels. For evaluating the performance of pixel-specific binary classification, we used various metrics including average precision (AP), mean area under the receiver operating characteristic (ROC) curve (AUC), and precision-recall curve (Pedregosa, et al., 2011). We also borrowed the metric of top accuracy from the community of protein contact map prediction (Kandathil, et al., 2019; Wang, et al., 2017). Specifically, we selected a set of pixels with the top N predicted scores and calculated the percentage (accuracy) of ground-truth pixels that are found in the set. If N is not given a specific number in downstream analysis, topN means that we select the top number of pixels that is equal to the ground-truth number of positive pixels for each chromosome.

## 3 Results

### 3.1 Hyperparameter selection

The testing results of hyperparameter tuning on chromosome 1 are shown in Table S4. The training data were generated from chromosomes 3 to X, while the validation data were extracted from chromosome 2. From the learned models with the positive weight 3, we observed that different batch sizes do affect validation loss and batch size 16 is a better selection for either one of the two normalization methods (instance or batch). After we fixed the batch size to 16, we found that compared with 0.01 and 0.0001 using the initial learning rate 0.001 achieved a smaller validation loss. Since batch sizes matter and instance normalizations are not related to batch size, we applied batch normalization for all convolutional layers in our network. We extended the number of residual blocks to 40; each block has its dilation value for dilated convolutions (Table S4). After increasing the hidden dimension to 128, we obtained the final optimal combination of hyperparameters that achieved the lowest validation loss: the batch size 16, the learning rate 0.001 with automatically reducing, the kernel size 3, and the positive weight 1. The training and validation loss curves that we obtained with the optimal hyperparameters are shown in Fig. S3. We did not observe overfitting; instead, we obtained an almost perfect loss curve for validation. To train the other 22 models for blindly testing on chromosomes from 2 to X, we used the same hyperparameters as we obtained from the testing on chromosome 1.

### 3.2 DeepChIA-PET(Hi-C) significantly outperforms Peakachu

To make a fair comparison with Peakachu, we trained DeepChIA-PET with only Hi-C data as input. The hyperparameters for training DeepChIA-PET(Hi-C) were the same as the ones we obtained in the hyperparameter-tuning process. The main difference is that the input channel is changed from three to one. We also made sure to train DeepChIA-PET(Hi-C) and Peakachu using the same input and ground-truth data. We used “score_chromosome” in Peakachu (Salameh, et al., 2020) to predict interaction probability per pixel for a chromosome. The comparison between Peakachu and DeepChIA-PET(Hi-C) is to directly evaluate their abilities in identifying pixel-specific ChIA-PET interactions.

We tested two chromosomes 1 and 2, and the comparison results are shown in Figures 2 and S4, respectively. With the increase of the number of negative pixels, the AP values keep reducing for both DeepChIA-PET(Hi-C) and Peakachu, whereas the AUCs have not been affected (Fig. 2A). Notably, DeepChIA-PET(Hi-C) achieved higher APs than Peakachu and almost perfect AUC (0.995) compared with 0.753 for Peakachu. In addition, the PR curves shown in Fig. 2B for neg=20pos also indicate that DeepChIA-PET(Hi-C) significantly outperforms Peakachu. Moreover, we compared top accuracy scores for the two methods (Fig. 2C), revealing that DeepChIA-PET(Hi-C) can successfully recover or pinpoint many more ground-truth pixels than Peakachu. We can draw the same conclusions from Fig. S4 for blindly testing on chromosome 2.

**Fig. 2.**
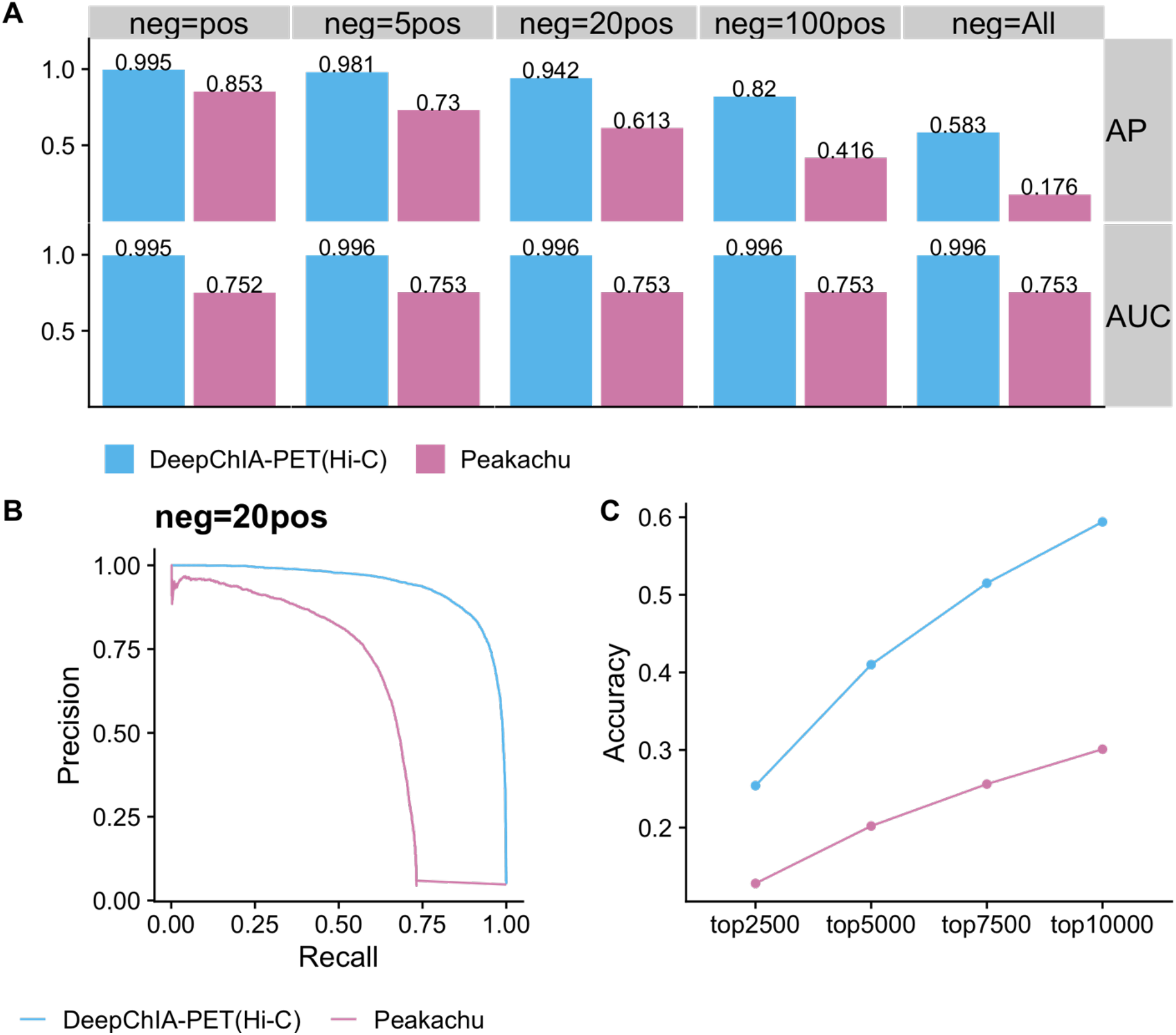
DeepChIA-PET(Hi-C) outperforms Peakachu for blindly testing CTCF ChIA-PET on chromosome 1 in GM12878. (A) The AP and ROC-AUC scores in terms of different numbers of negative pixels when evaluating. (B) Precision-recall curve for the number of negative pixels equaling 20 times the number of positive pixels. (C) The topN accuracy where N equals four different values.

### 3.3 ChIP-seq data improve the performance of DeepChIA-PET

We next investigated the contribution of ChIP-seq data in DeepChIA-PET. We trained four more DeepChIA-PET models with only Hi-C as input, with only ChIP-seq as input, and with both ChIP-seq and Hi-C as input for blindly testing on chromosomes 1 and 2 and the evaluation results are shown in Figures S5 and 3, respectively. As in the comparisons between DeepChIA-PET(Hi-C) and Peakachu, we calculated AP and AUC with different numbers of negative pixels, drew the PR curves for neg=20pos, and obtained the topN accuracy scores. From Fig. 3, we can conclude that (1) DeepChIA-PET with both Hi-C and ChIP-seq as input consistently outperforms the other two cases in all the metrics; (2) the performance of DeepChIA-PET with Hi-C as input, or DeepChIA-PET(Hi-C), is much more close to DeepChIA-PET(Hi-C and ChIA-PET) than DeepChIA-PET with only ChIP-seq as input, or DeepChIA-PET(ChIP-seq); and (3) adding ChIP-seq along with Hi-C does significantly improve the performance of our classifier. The same conclusions can be drawn when blindly testing on chromosome 1 (Fig. S5).

**Fig. 3.**
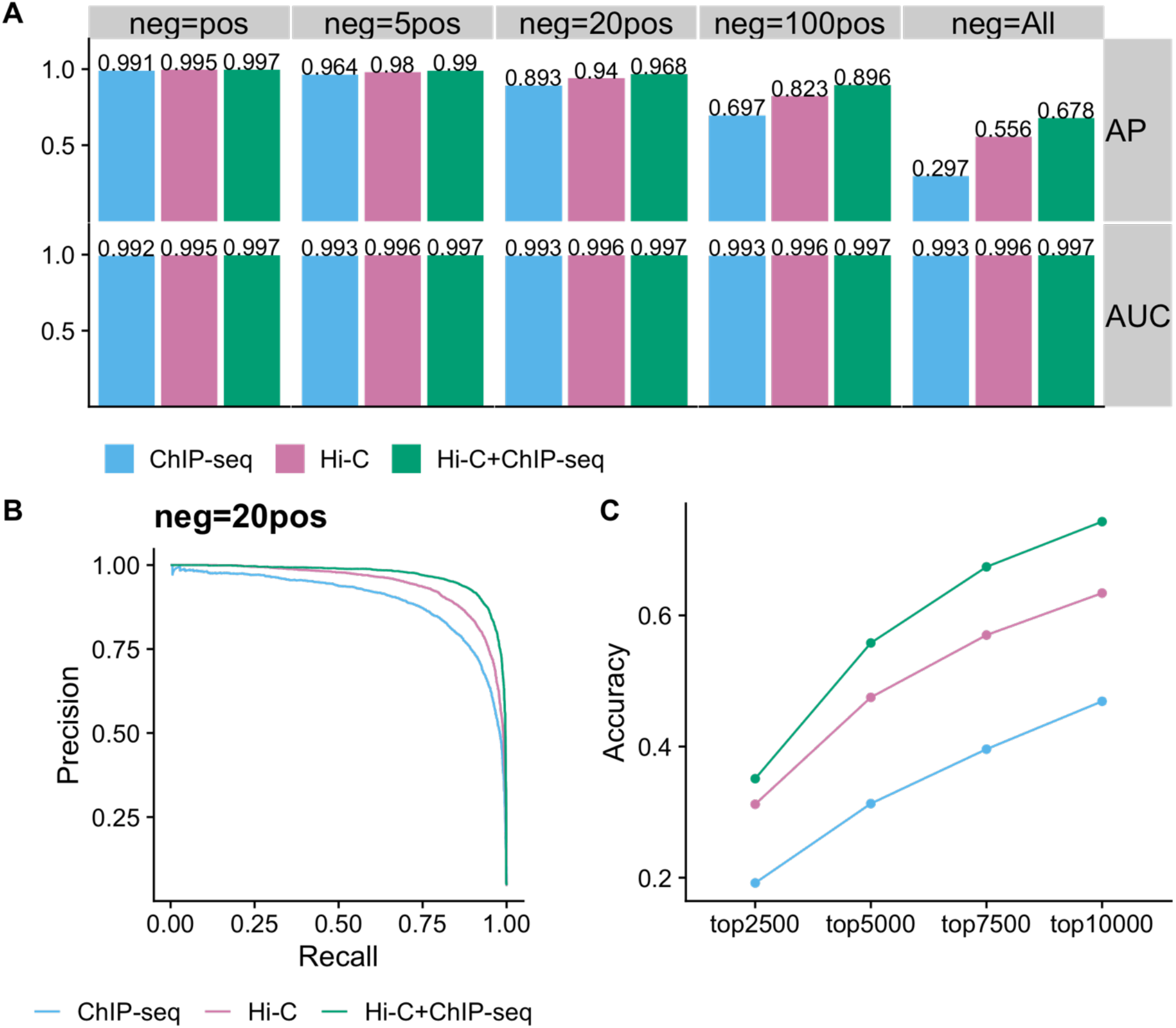
Ablation studies of the contribution of ChIP-seq. The evaluations for DeepChIA-PET with only Hi-C, only ChIP-seq, and both as input were conducted on chromosome 2 for CTCF ChIA-PET in GM12878. (A) The AP and ROC-AUC scores for different numbers of negative pixels. (B) Precision-recall curve for neg=20pos. (C) The topN accuracy where N equals four different values.

### 3.4 Overall performance on CTCF ChIA-PET in GM12878

DeepChIA-PET achieved state-of-the-art performance on all testing chromosomes from chromosome 1 to the X-chromosome in GM12878 (Fig. 4A). Specifically, we achieve almost perfect AUC (mean ≥ 0.997) for all five different numbers of negative pixels, and the higher AP values also indicate that DeepChIA-PET can successfully identify positive pixels among an imbalanced pool that is heavily occupied by negatives.

**Fig. 4.**
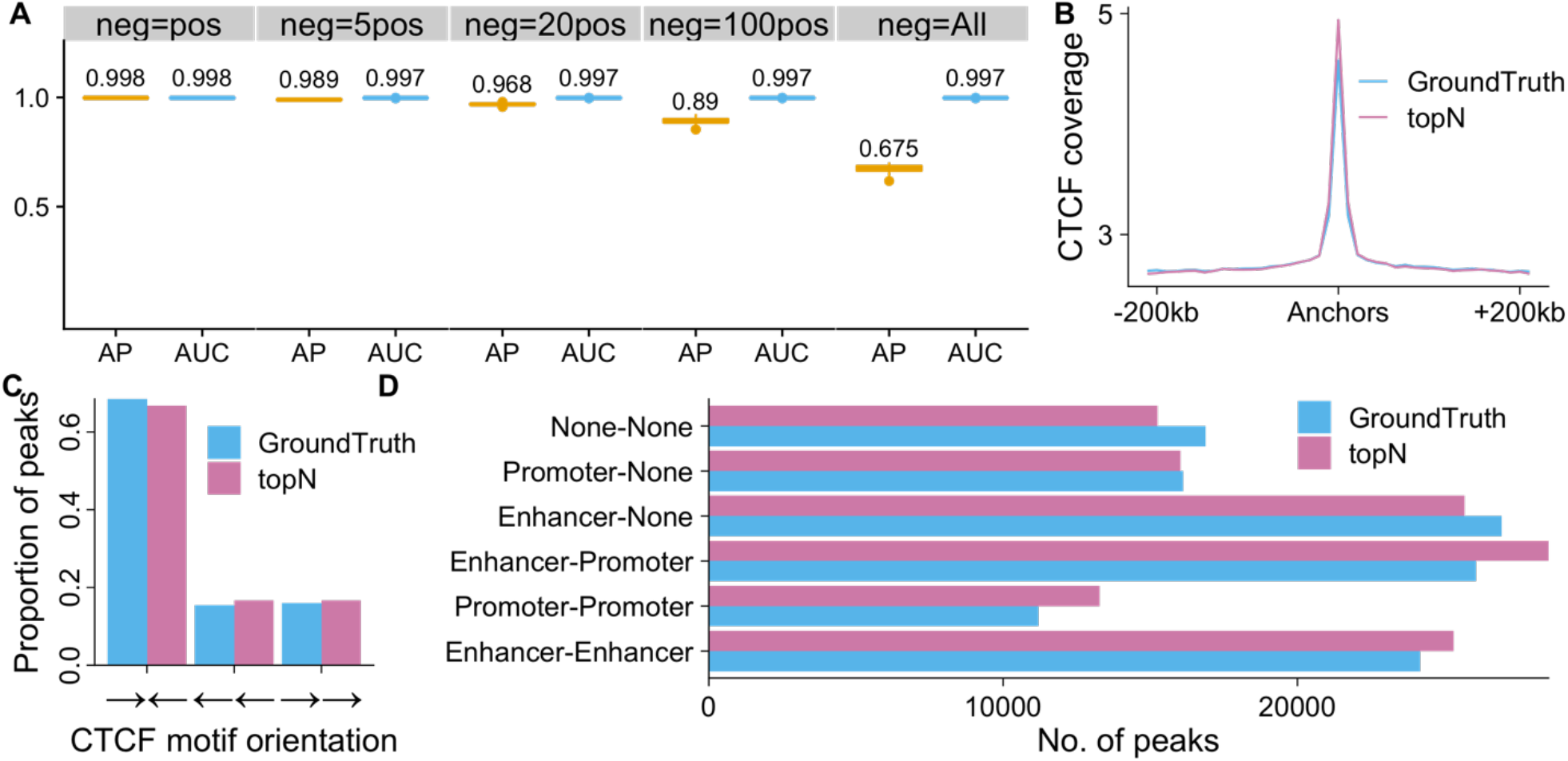
Overall performance of DeepChIA-PET for predicting CTCF ChIA-PET interactions on all testing chromosomes in GM12878. (A) The boxplots of APs and ROC-AUCs for all testing chromosomes (mean values are added above each boxplot). (B) The average CTCF coverages surrounding anchors from ground truth and our predicted topN pixels. (C) CTCF motif orientation analysis at anchors from ground truth and our predicted topN pixels. (D) ChIA-PET interactions between promoters and enhancers.

Next, we focused on our predicted topN pixels to find out if these pixels have similar transcriptional properties with ground-truth pixels. We first found that the genomic regions surrounding anchors from both topN and ground truth are enriched with CTCF (Fig. 4B), which is what we expected for CTCF ChIA-PET interactions. We then found CTCF motif orientation using MotifFinder in Juicer (Durand, et al., 2016) and observed that the pairs of CTCF motifs that anchor most of topN and ground-truth pixels (> 60%) are in the convergent orientation (Fig. 4C) and our topN pixels have very similar CTCF motif orientation (convergent and tandem) with ground truth.

Moreover, we assigned regulatory elements (promoters, enhancers, or none for not finding any promoter or enhancer) to each anchor at topN and ground-truth loci and counted the numbers of pixels that belong to each of the six different regulatory element combinations. The promoter and enhancer loci of GM12878 were extracted from ChromHMM segmentations in ENCODE with 10 states. We found that most of the pixels in both topN and ground truth are related to at least one of the well-known regulatory elements (Fig. 4D). Particularly, our topN set contains more significant interactions than ground truth for the three regulatory element combinations (enhancer-enhancer, promoter-promoter, and enhancer-promoter).

We showed two specific examples of our predicted CTCF ChIA-PET interactions on chromosomes 1 (Fig. 5) and 10 (Fig. S6) in GM12878. We observed several TADs in the heat map of KR-normalized Hi-C. We also found that the insulator-binding protein CTCF coverages are enriched not only in surrounding TAD boundary regions but also at some non-boundary loci. Moreover, we found that the anchors of ground-truth ChIA-PET interactions are usually located between the genomic regions that are enriched for the binding of CTCF, and the heat map of our predicted CTCF ChIA-PET is so accurate that it looks like a mirror image of the ground-truth heat map.

**Fig. 5.**
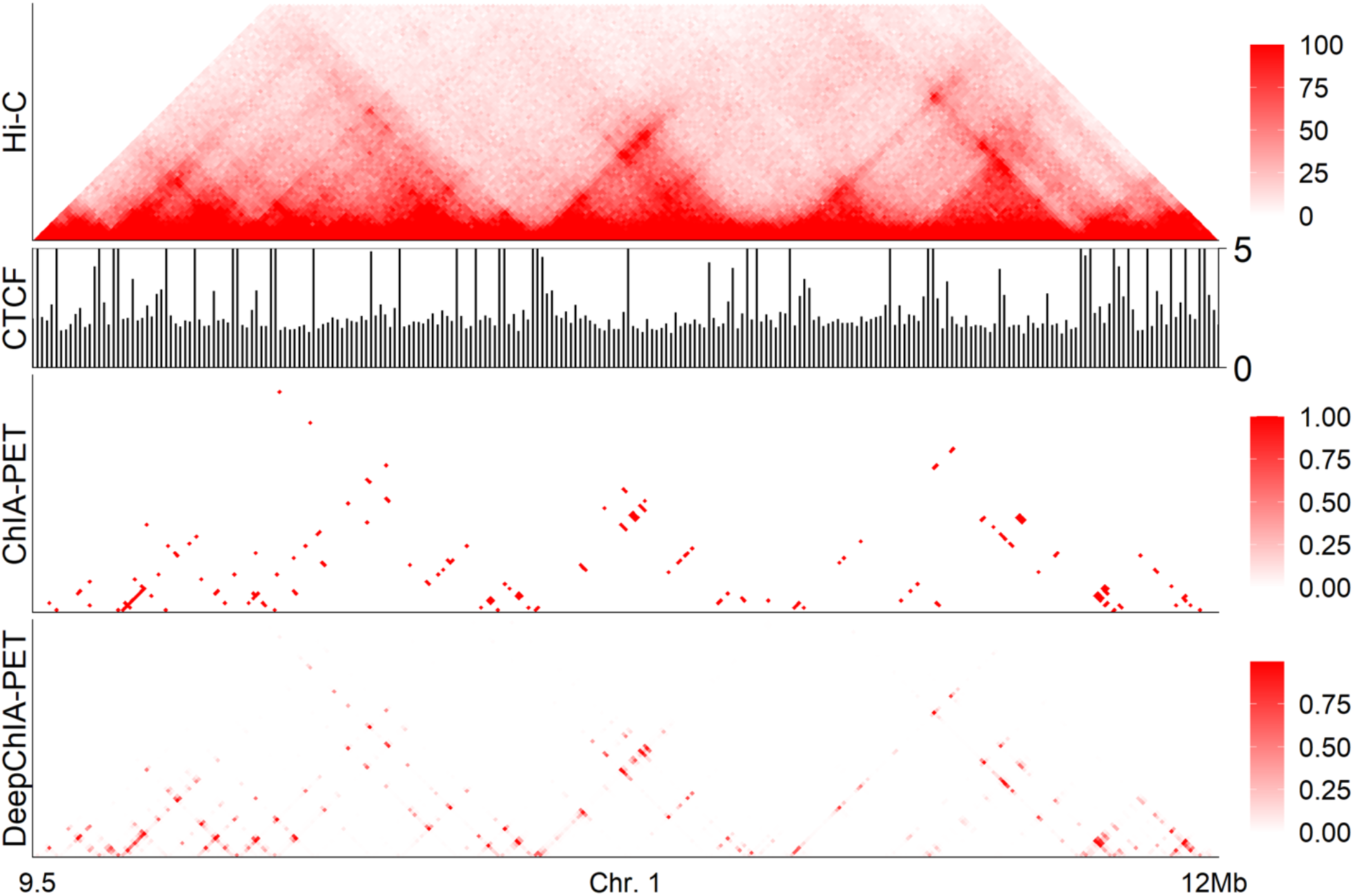
A specific example of our predictions for CTCF ChIA-PET interactions on chromosome 1 in GM12878. From top to bottom: the KR-normalized Hi-C, the CTCF coverage at 10-kb resolution, the heat map for ground-truth ChIA-PET interactions, and the heat map for our predicted ChIA-PET interactions.

### 3.5 Overall performance on CTCF ChIA-PET in HeLa

We next evaluated DeepChIA-PET on CTCF ChIA-PET in a different cell type HeLa. We first checked the similarities of ground-truth CTCF ChIA-PET between GM12878 and HeLa. We found that the number of HeLa loops is less than half of the number of GM12878 loops and ∼65.5% HeLa ChIA-PET loops were found in GM12878 (Fig. 6A), indicating that HeLa ChIA-PET interactions are sparser than GM12878.

**Fig. 6.**
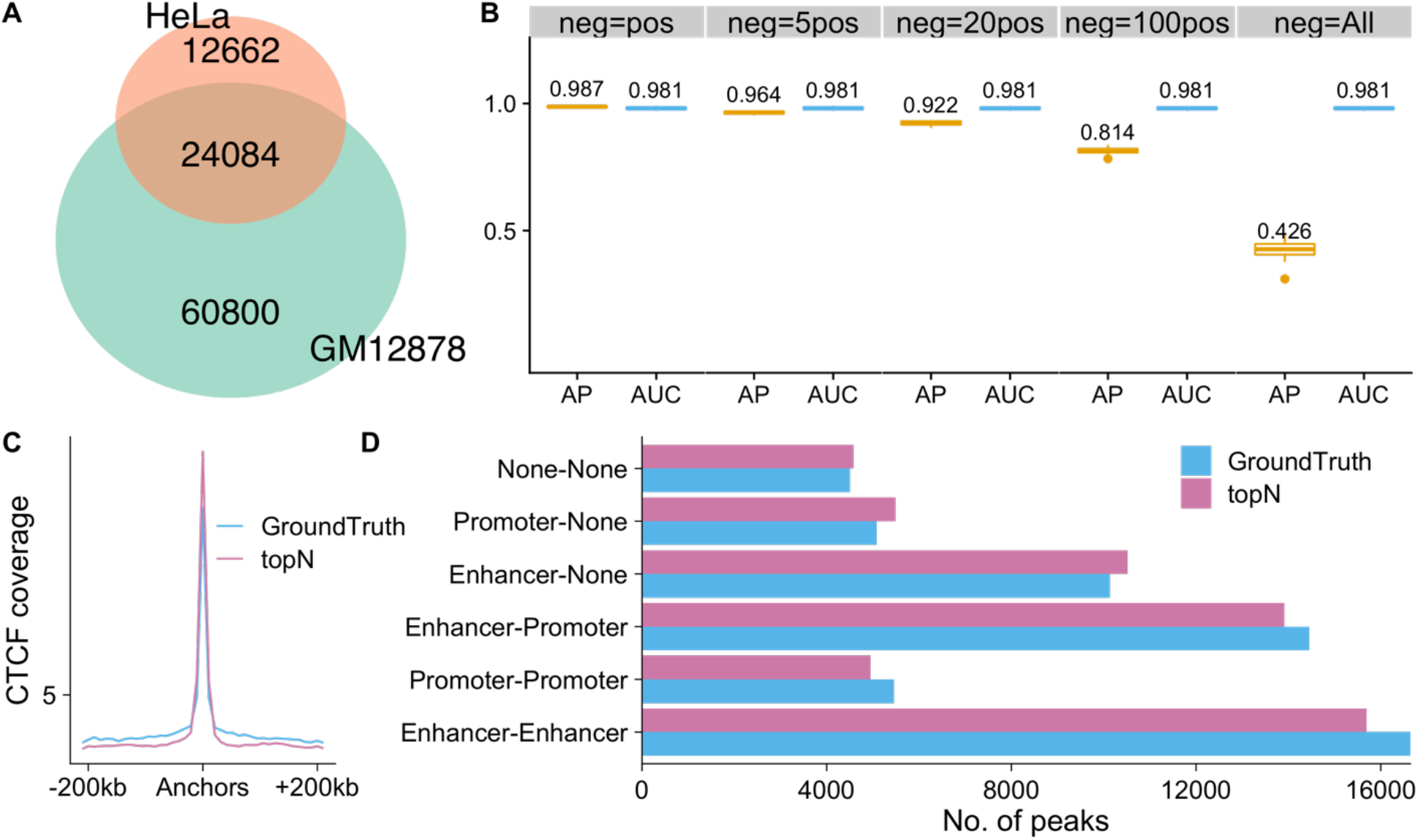
Overall performance of DeepChIA-PET for CTCF ChIA-PET predictions in a different cell type HeLa. (A) Overlap in ground-truth ChIA-PET interactions between GM12878 and HeLa. (B) The boxplots of APs and ROC-AUCs for all testing chromosomes (mean values are added above each boxplot). (C) The average CTCF coverages surrounding anchors from ground truth and our predicted topN pixels. (D) ChIA-PET interactions between promoters and enhancers.

The overall performance of DeepChIA-PET for predicting CTCF ChIA-PET in HeLa is shown in Fig. 6B. Compared with the performance in GM12878, the AUC values have a slight decrease from 0.997 to 0.981, and the AP values have a noticeable drop when using all negative pixels (neg=All), which may result from the massively increasing of the number of negative pixels. The anchors from ground truth and the topN set in HeLa are enriched for CTCF (Fig. 6C), and the two sets (topN and ground truth) have very similar loop distributions for the six different regulatory element combinations (Fig. 6D). As in GM12878, the promoter and enhancer loci of HeLa were extracted from ChromHMM segmentations in ENCODE.

A specific example of our predictions on chromosome 6 in HeLa is shown in Fig. 7, we can make the same conclusions as we made from Figures 5 and S6. In summary, DeepChIA-PET that was trained with CTCF ChIA-PET in GM12878 can accurately predict CTCF ChIA-PET in HeLa.

**Fig. 7.**
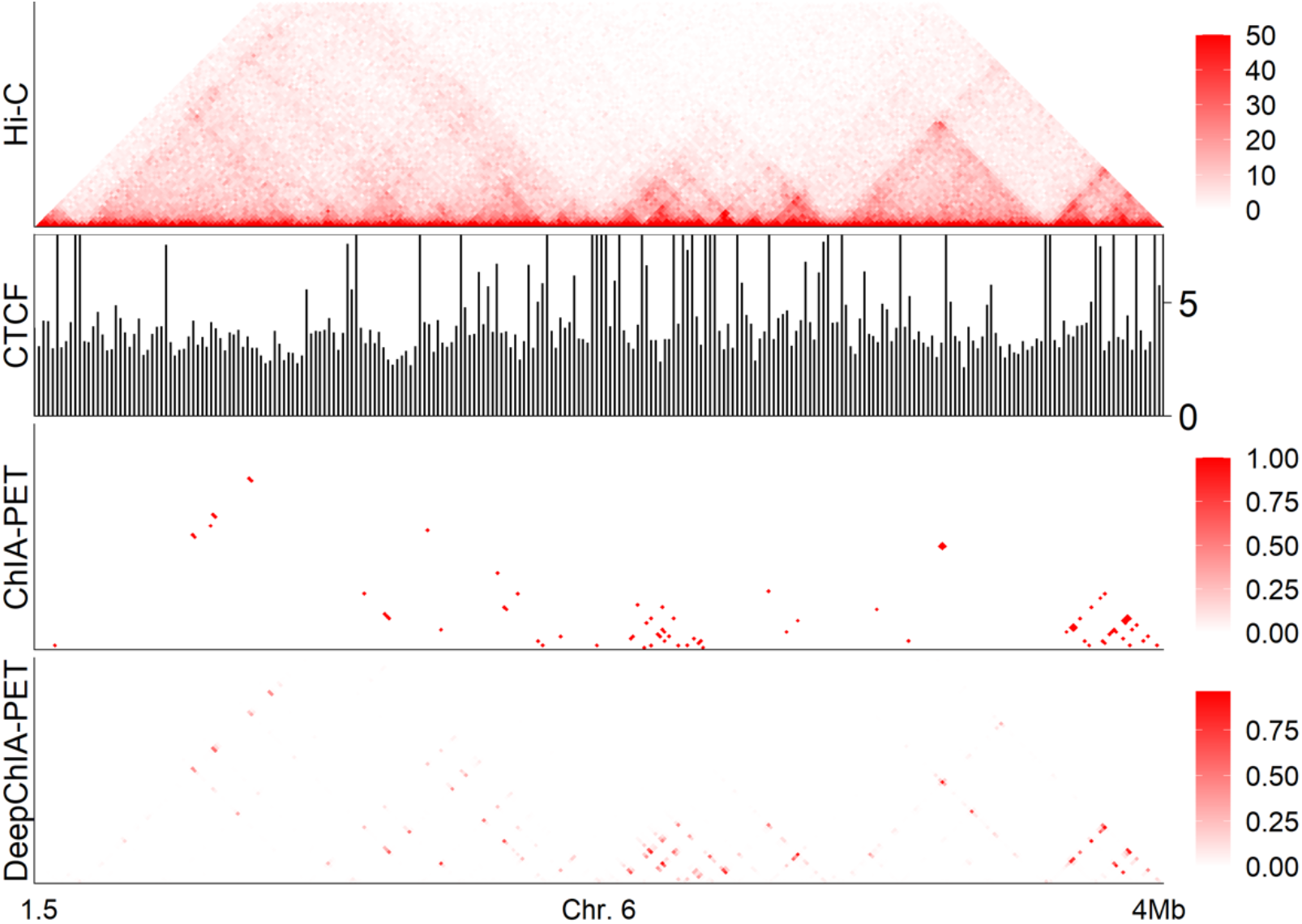
A specific example of our predictions for CTCF ChIA-PET on chromosome 6 in HeLa. From top to bottom: KR-normalized Hi-C, CTCF coverage at 10-kb resolution, ground-truth ChIA-PET, and our predicted ChIA-PET.

### 3.6 Overall performance on RNAPII ChIA-PET in GM12878 and HeLa

Since ChIA-PET experiments can be conducted on different DNA-binding proteins, we used DeepChIA-PET to predict RNAPII ChIA-PET in GM12878 and HeLa. We first compared the similarities between CTCF ChIA-PET of GM12878 and RNAPII ChIA-PET in GM12878 and HeLa (Fig. 8A and B) and found that RNAPII ChIA-PET does not share most of its loops with CTCF ChIA-PET even through the cell types are the same.

**Fig. 8.**
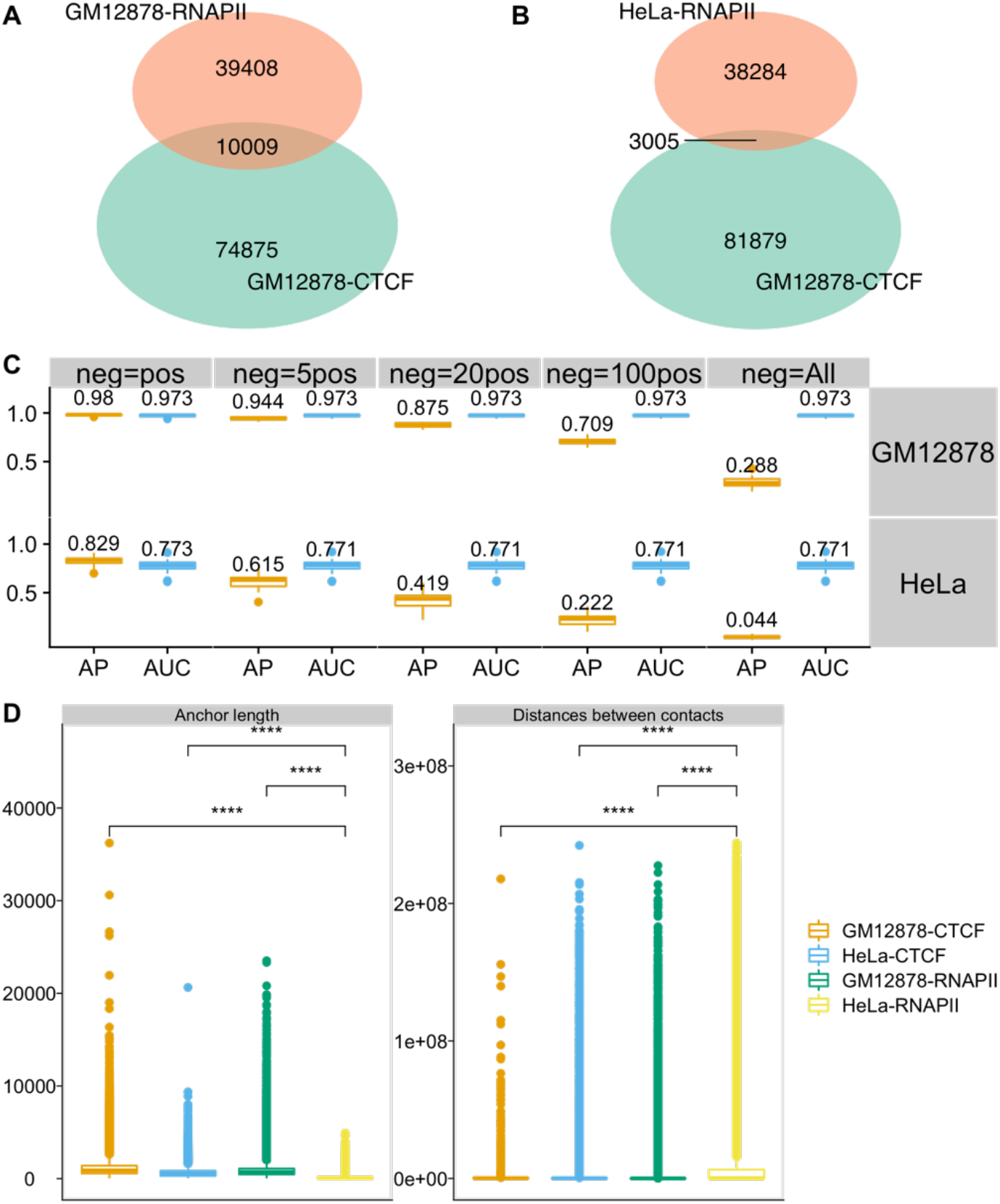
Overall performance of DeepChIA-PET on predicting RNAPII ChIA-PET in GM12878 and HeLa. The overlaps in ground-truth pixel-wise ChIA-PET interactions between CTCF GM12878 and RNAPII GM12878 are shown in (A), and between CTCF GM12878 and RNAPII HeLa are shown in (B). (C) The boxplots of APs and ROC-AUCs for all testing chromosomes in both GM1278 and HeLa. (D) RNAPII ChIA-PET interactions for HeLa are significantly different from either the RNAPII ChIA-PET in GM12878 or the CTCF ChIA-PET in both GM12878 and HeLa by having shorter anchor lengths and longer genomic distances between interactions. Mean comparisons were conducted with student’s t-Test (****: p <= 0.0001).

The genome-wide AP and AUC values of DeepChIA-PET are shown in Fig. 8C. For predicting RNAPII ChIA-PET of GM12878, our tool achieved a higher AUC of 0.973, and the AP performs well except when we used all negative pixels (neg=All) for evaluation. However, for predicting RNAPII ChIA-PET of HeLa, we achieved an AUC of 0.771, and the AP values keep reducing with the increase of negative pixels used for evaluation. The reason that DeepChIA-PET did not perform well in predicting RNAPII ChIA-PET of HeLa may be that RNAPII ChIA-PET loops of HeLa have shorter anchor lengths and longer genomic distances (Fig. 8D), which is significantly different from what we found from CTCF ChIA-PET of GM12878. Together, DeepChIA-PET can accurately predict RNAPII ChIA-PET of GM12878, but not perform that well in RNAPII ChIA-PET of HeLa.

### 3.7 Overall performance on RAD21 ChIA-PET in GM12878 and K562 from different experiments

The CTCF and RNAPII ChIA-PET data that we used for evaluation are from the same study of (Tang, et al., 2015). In this section, we evaluated DeepChIA-PET in predicting RAD21 ChIA-PET interactions, which were obtained from different experiments (Heidari, et al., 2014).

As before, we first compared the similarities of ChIA-PET loops between CTCF of GM12878 and RAD21 of GM12878, and between CTCF of GM12878 and RAD21 of K562. We found ∼87.1% RAD21 loops of GM12878 are shared with CTCF loops of GM12878 (Fig. 9A) and ∼48.6% RAD21 loops of K562 are also detected in CTCF loops of GM12878 (Fig. 9B). Therefore, the two RAD21 ChIA-PET datasets are more similar to CTCF ChIA-PET of GM12878 than between the two RANPII ChIA-PET loop sets.

**Fig. 9.**
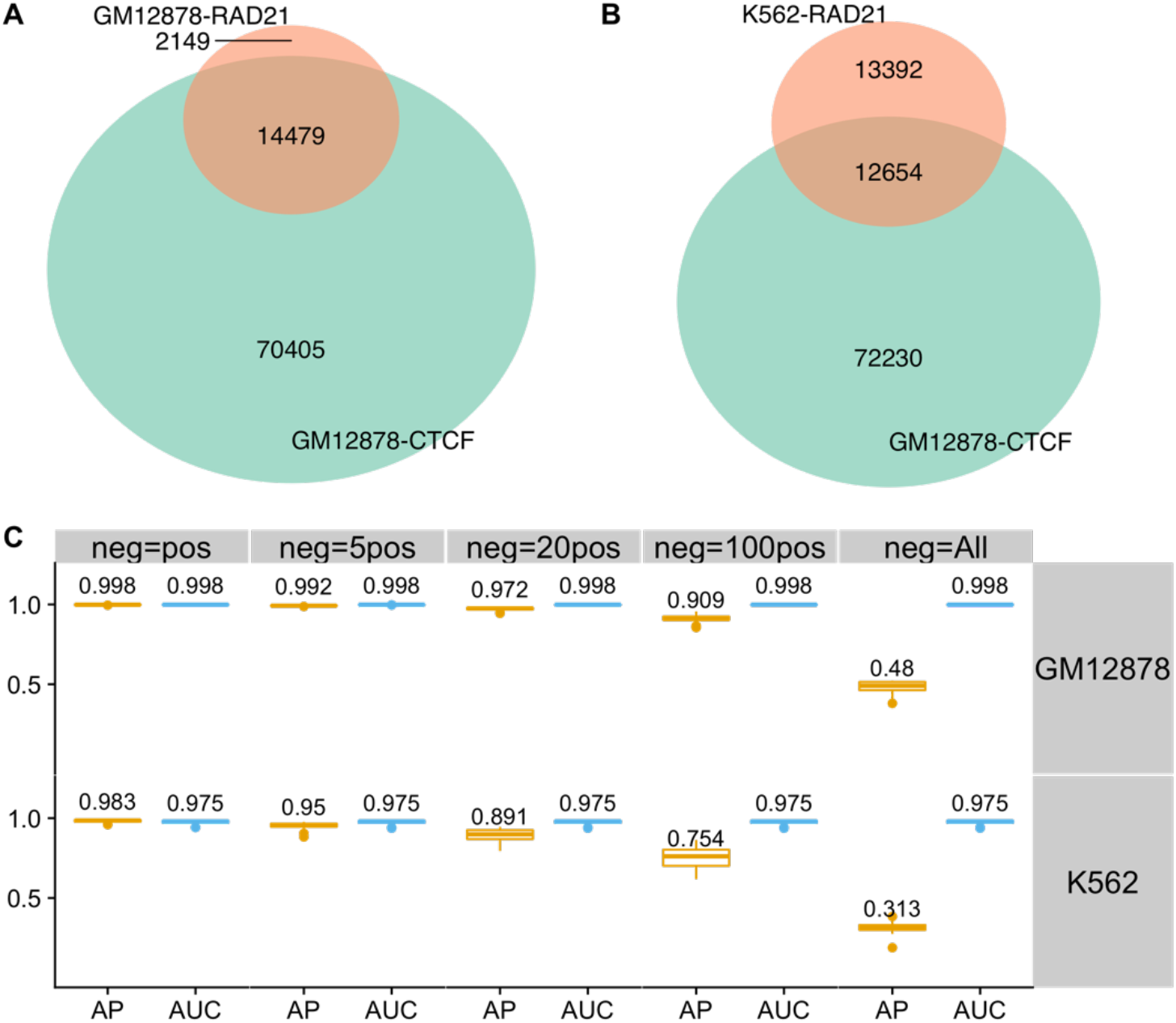
Overall performance on predicting RAD21 ChIA-PET in both GM12878 and K562. The overlaps in ground-truth pixel-wise ChIA-PET interactions between CTCF GM12878 and RAD21 GM12878 are shown in (A), and between CTCF GM12878 and RAD21 K562 are shown in (B). (C) The boxplots of APs and ROC-AUCs for all testing chromosomes in both GM1278 and K562. Note: since the Hi-C matrix for chromosome 9 in K562 is empty, the evaluation results shown in (C) for K562 do not include the testing scores for chromosome 9.

We reported AP and ROC-AUC scores of DeepChIA-PET for predicting RAD21 ChIA-PET of GM12878 and RAD21 ChIA-PET of K562 (Fig. 9C). DeepChIA-PET achieved almost perfect ROC-AUCs (0.998 for GM12878 and 0.975 for K562) and also obtained high AP values (0.972 for GM12878 and 0.891 for K562) when neg=20pos. We also reported different topN accuracy scores obtained on all testing chromosomes from chromosome 1 to the X-chromosome (Fig. S7) and found that the more top predicted pixels that we considered for evaluations, the higher topN accuracy we obtained, which can be used as a guide when deciding on the number of top predicted interactions to be used in applications.

### 3.8 Comparison between CTCF ChIA-PET and Hi-C peaks

The anchors of Hi-C peaks/loops called on Hi-C by HiCCUPS have been typically found at TAD boundaries and CTCF binding sites (Rao, et al., 2014). We detected 8609 Hi-C peaks on Hi-C contact matrices of GM12878 at 10-kb resolution. We reported the overlaps between Hi-C peaks and ground-truth CTCF ChIA-PET interactions of GM12878 (Fig. 10A), and between Hi-C peaks and our topN predicted set (Fig. 10B). It is observed that most of the Hi-C peaks can also be found in the two CTCF ChIA-PET loop sets: 83.6% Hi-C peaks are shared with ground-truth ChIA-PET, whereas 87.7% Hi-C peaks are also found in topN, indicating that DeepChIA-PET has the ability to select Hi-C peaks as potential ChIA-PET loops.

**Fig. 10.**
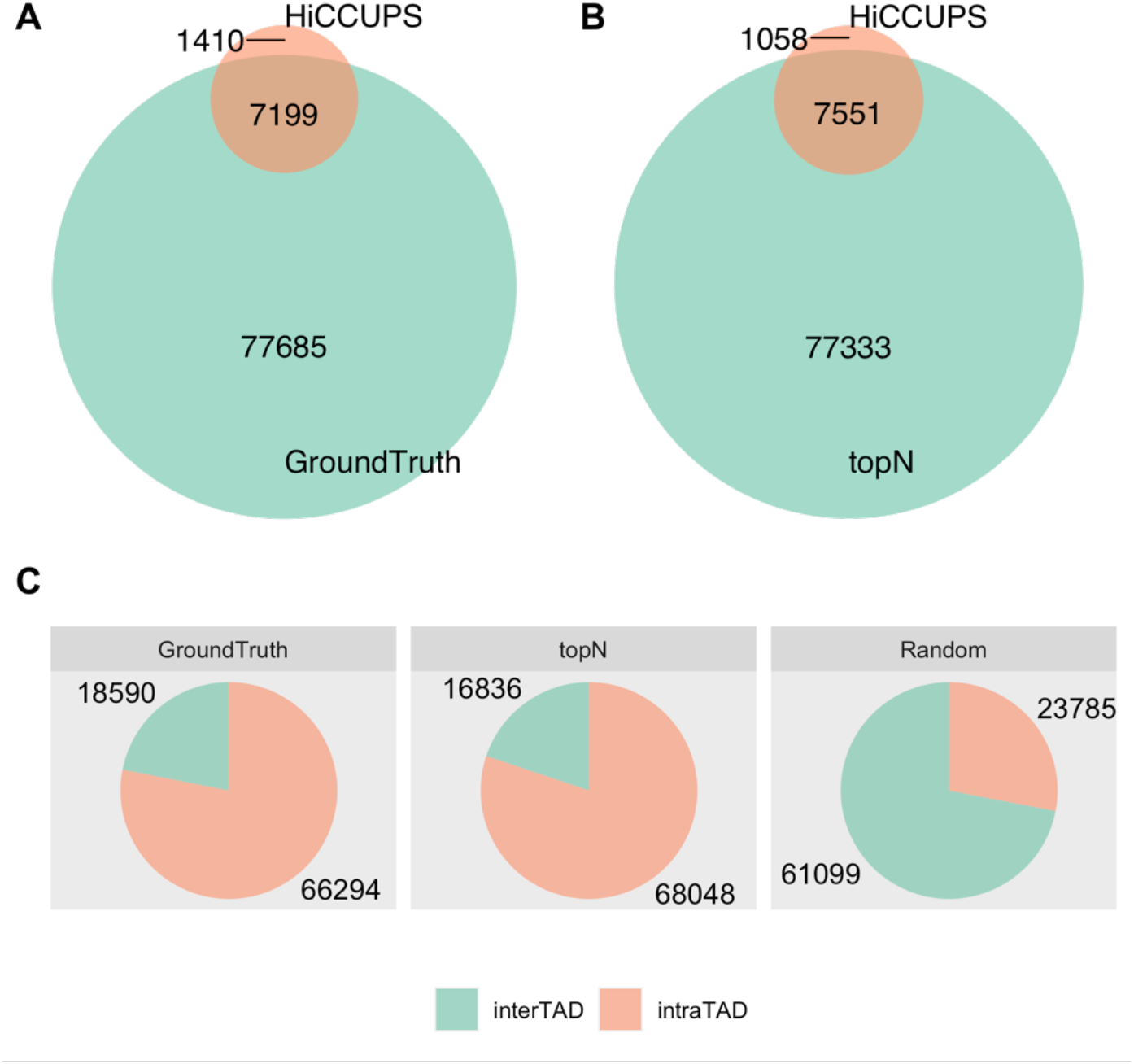
Genome-wide overlaps between Hi-C peaks called by HiCCUPS and ground-truth CTCF ChIA-PET interactions (A), and between Hi-C peaks called by HiCCUPS and our predicted topN CTCF ChIA-PET interactions (B) in GM12878. When counting the accordant pixels, we allow ± 1 bin mismatch. (C) Compared with randomly selected pixels, most of the ground-truth and our predicted topN CTCF ChIA-PET interactions are found within TADs.

In addition, we explored the relationships between CTCF ChIA-PET interactions and TADs. We found that compared with 28% of randomly selected pixels, more than 78% of ground-truth and 80% of predicted topN CTCF ChIA-PET interactions are found within TADs (Fig. 10C).

## Conclusions

In this study, we present DeepChIA-PET, a supervised deep-learning method for predicting ChIA-PET from Hi-C and ChIP-seq data. Our evaluation results indicate that DeepChIA-PET with only Hi-C as input significantly outperforms Peakachu. The ablation studies prove that ChIP-seq data as input contribute to the classification task of DeepChIA-PET. For predicting CTCF ChIA-PET in GM12878 and HeLa, DeepChIA-PET can achieve ROC-AUCs of 0.997 and 0.973, and our predicted topN loops have very similar patterns of CTCF motif orientation with ground-truth ChIA-PET interactions in GM12878. The regulatory elements are widely found at most of the anchors from our topN loops, and the distributions of different regulatory element interactions are very similar to those from the ground truth. We also reported that DeepChIA-PET can be used to accurately predict non-CTCF ChIA-PET interactions, including RNAPII ChIA-PET of GM12878, RAD21 ChIA-PET of GM12878, and RAD21 ChIA-PET of K562, even though these ChIA-PET interactions are usually different from CTCF ChIA-PET of GM12878 in terms of the number of total and overlapping interactions. At last, we compared CTCF ChIA-PET interactions with Hi-C peaks and TADs and found that our topN loops have more common pixels with Hi-C peaks than ground truth and the number of our topN loops that are located within TADs is more than ground truth.

