## supplementary document for "DeepChIA-PET: Accurately predicting ChIA-PET from Hi-C and ChIP-seq with deep dilated networks"

### Supplementary Materials

**Table S1.** The source of Hi-C datasets.

| Cell type | Source |
|---|---|
| GM12878 | https://hicfiles.s3.amazonaws.com/hiseq/gm12878/in-situ/primary.hic |
| HeLa | https://hicfiles.s3.amazonaws.com/hiseq/hela/in-situ/combined.hic |
| K562 | https://hicfiles.s3.amazonaws.com/hiseq/k562/in-situ/combined.hic |

**Table S2.** The source of ChIP-seq datasets.

| ChIP-seq | Cell type | Source |
|---|---|---|
| CTCF | GM12878 | http://hgdownload.cse.ucsc.edu/goldenPath/hg19/encodeDCC/wgEncodeBroadHistone/wgEncodeBroadHistoneGm12878CtcfStdSig.bigWig |
| | HeLa | http://hgdownload.cse.ucsc.edu/goldenPath/hg19/encodeDCC/wgEncodeBroadHistone/wgEncodeBroadHistoneHelas3CtcfStdSig.bigWig |
| RNA PII | GM12878 | http://hgdownload.cse.ucsc.edu/goldenPath/hg19/encodeDCC/wgEncodeSydhTfbs/wgEncodeSydhTfbsGm12878Pol2IggmusSig.bigWig |
| | HeLa | http://hgdownload.cse.ucsc.edu/goldenPath/hg19/encodeDCC/wgEncodeSydhTfbs/wgEncodeSydhTfbsHelas3Pol2StdSig.bigWig |
| RAD 21 | GM12878 | http://hgdownload.cse.ucsc.edu/goldenPath/hg19/encodeDCC/wgEncodeSydhTfbs/wgEncodeSydhTfbsGm12878Rad21IggrabSig.bigWig |
| | K562 | http://hgdownload.cse.ucsc.edu/goldenPath/hg19/encodeDCC/wgEncodeSydhTfbs/wgEncodeSydhTfbsK562Rad21StdSig.bigWig |

**Table S3.** The source of ChIA-PET datasets.

| ChIA-PET | Source |
|---|---|
| GM12878-CTCF | GEO, GSM1872886 |
| GM12878-RNAPII | GEO, GSM1872887 |
| HeLa-CTCF | GEO, GSM1872888 |
| HeLa-RANPII | GEO, GSM1872889 |
| GM12878-RAD21 | Table S1 from Heidari et al (2014) Genome research. |
| K562-RAD21 | |

**Table S4.** Results for hyperparameter tuning. The batch size for validation data was set to one for all models. Learning rates followed by "(↓)" indicate that we reduced the learning rate by a factor of 0.1 when validation loss stops improving. Positive weights followed by a * mean that we only used this positive weight when calculating training loss; if there is no * we used the positive weight for calculating both training and validation losses. Therefore, we can compare validation loss between models with the same positive weight without having * or models with positive weights followed by a *. All models are trained for blindly testing on chromosome 1, which means we extract training data from chromosome 3 up to X and validation data from chromosome 2.

| Mid | Batch size | Learning rate | Kernel size | Norm | No. of residual blocks | Dilation for residual blocks | Hidden dimension | Positive weight | Validation loss |
|---|---|---|---|---|---|---|---|---|---|
| 1 | 4 | 0.001 | 5 | Instance | 6 | [1,2,1,4,1,1] | 64 | 3 | 0.00932 |
| | 8 | 0.001 | 5 | Instance | 6 | | 64 | 3 | 0.00913 |
| | 16 | 0.001 | 5 | Instance | 6 | | 64 | 3 | 0.0091 |
| | 32 | 0.001 | 5 | Instance | 6 | | 64 | 3 | 0.01077 |
| | 16 | 0.01 | 5 | Instance | 6 | | 64 | 3 | 0.01005 |
| | 16 | 0.0001 | 5 | Instance | 6 | | 64 | 3 | 0.09291 |
| 2 | 8 | 0.001 | 5 | Batch | 6 | | 64 | 3 | 0.00939 |
| | 16 | 0.001 | 5 | Batch | 6 | | 64 | 3 | 0.00932 |
| 3 | 16 | 0.001 | 5 | Instance | 10 | [1,2,1,4,1,8,1,16,1,1] | 64 | 3 | 0.00904 |
| 4 | 16 | 0.001 | 5 | Instance | 14 | [1,2,1,4,1,8, 1,16,1,32,1,64,1,1] | 64 | 3 | 0.00901 |
| 5 | 16 | 0.001 (↓) | 5 | Batch | 14 | | 64 | 3 | 0.0088 |
| | 32 | 0.001 (↓) | 5 | Batch | 14 | | 64 | 3 | 0.00909 |
| | 64 | 0.001 (↓) | 5 | Batch | 14 | | 64 | 3 | 0.0099 |
| 6 | 16 | 0.001 (↓) | 5 | Batch | 20 | [1,1,2,1,1,4,1,1,8,1, 1,16,1,1,32,1,1,64,1,1] | 64 | 3 | 0.00897 |
| 7 | 16 | 0.001 (↓) | 3 | Batch | 14 | [1,2,1,4,1,8, 1,16,1,32,1,64,1,1] | 64 | 3 | 0.00875 |
| | 16 | 0.001 (↓) | 3 | Batch | 14 | | 64 | 1 | 0.0046 |
| 8 | 16 | 0.001 (↓) | 3 | Batch | 20 | [1,1,2,1,1,4,1,1,8,1, 1,16,1,1,32,1,1,64,1,1] | 64 | 3 | 0.00883 |
| | 16 | 0.001 (↓) | 3 | Batch | 20 | | 64 | 6 | 0.01275 |
| | 16 | 0.001 (↓) | 3 | Batch | 20 | | 64 | 12 | 0.01891 |
| 9 | 16 | 0.001 (↓) | 3 | Batch | 20 | | 128 | 6 | 0.01324 |
| 10 | 16 | 0.001 (↓) | 3 | Batch | 40 | [1,1,2,1,1,4,1,1,8, 1,1,16,1,1,32,1,1,64, 1,1,1,1,1,1,1,1,1,1,1,1, 1,1,1,1,1,1,1,1,1,1] | 64 | 6 | 0.01317 |
| 11 | 16 | 0.001 (↓) | 3 | Batch | 40 | | 128 | 6 | 0.01319 |
| 12 | **16** | **0.001 (↓)** | **3** | **Batch** | **40** | | **128** | **1*** | **0.00458** |
| | 16 | 0.001 (↓) | 3 | Batch | 40 | | 128 | 3* | 0.00479 |
| | 16 | 0.001 (↓) | 3 | Batch | 40 | | 128 | 6* | 0.00625 |
| | 16 | 0.001 (↓) | 3 | Batch | 40 | | 128 | 9* | 0.00552 |
| 13 | 16 | 0.001 (↓) | 5 | Batch | 40 | | 128 | 1* | 0.0047 |
| | 16 | 0.001 (↓) | 5 | Batch | 40 | | 128 | 3* | 0.0049 |
| | 16 | 0.001 (↓) | 5 | Batch | 40 | | 128 | 6* | 0.00507 |
| | 16 | 0.001 (↓) | 5 | Batch | 40 | | 128 | 9* | 0.00601 |

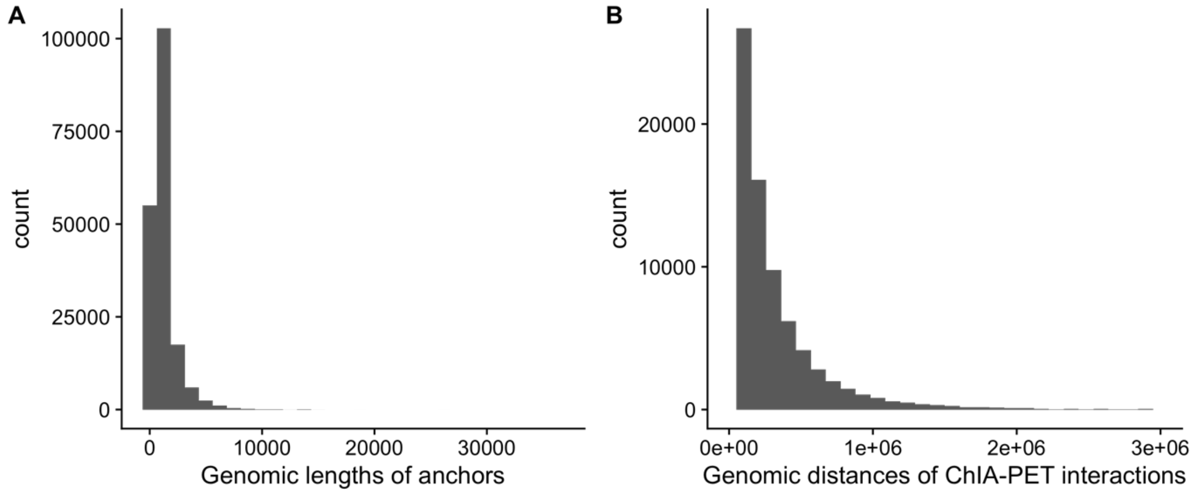


**Figure S1.** For CTCF ChIA-PET interactions in GM12878, we find (A) almost all anchors' genomic lengths are less than 10 kb, and (B) almost all ChIA-PET interactions are within 2 Mb.

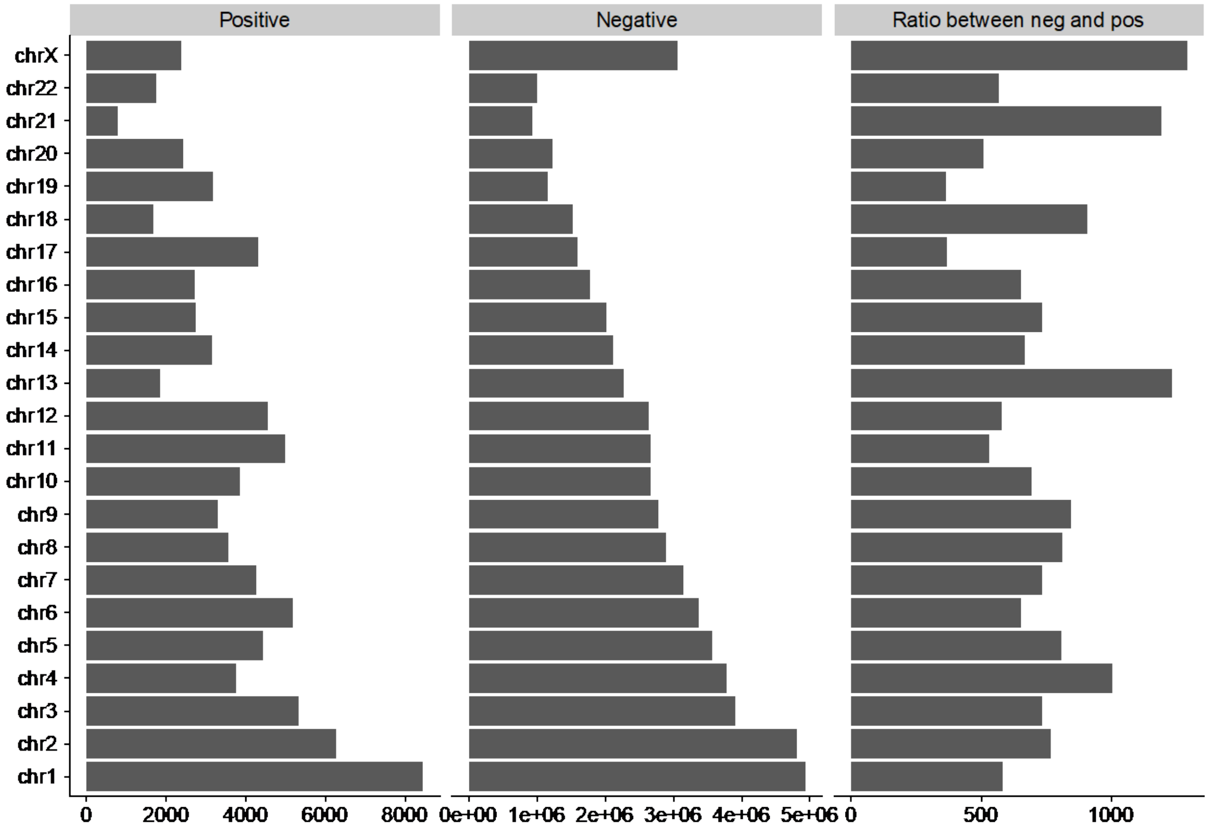


**Figure S2.** Number of positive and negative pixels for each chromosome and the ratios between negative and positive at 10-kb resolution. The range of genomic distances we cared about is from 2 to 200 bins, that is, 20 kb to 2 Mb. The ChIA-PET data are from CTCF of GM12878.

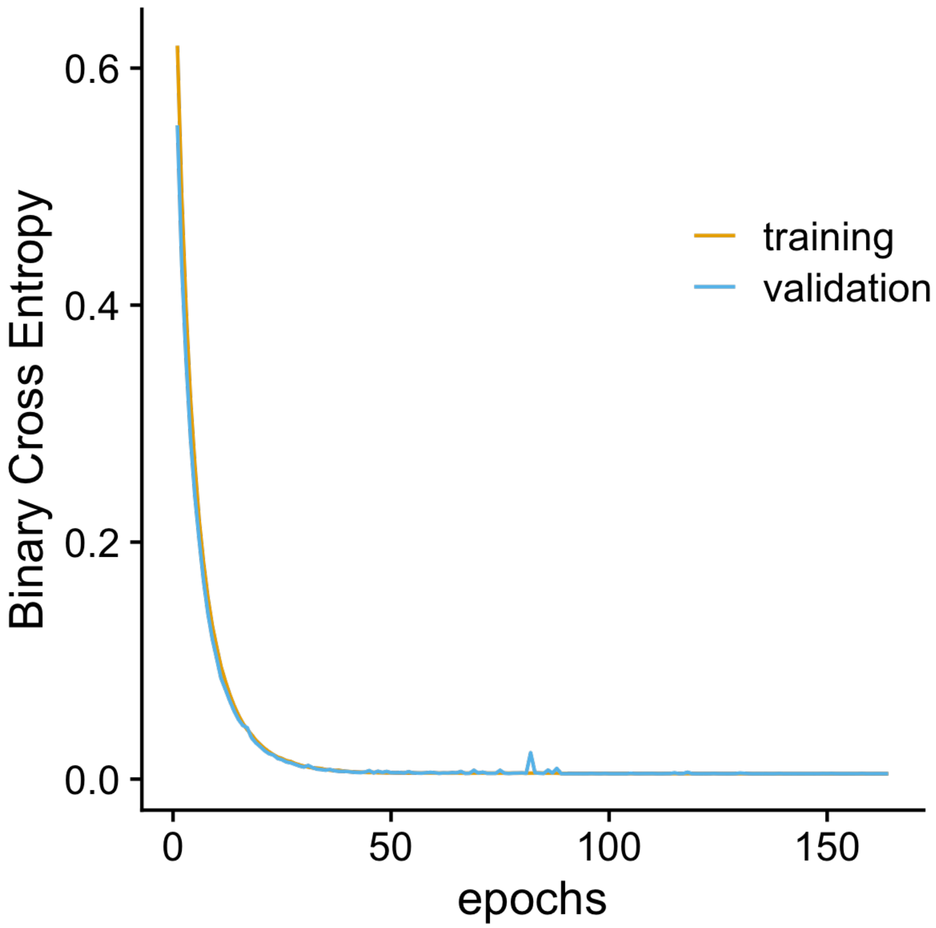


**Figure S3.** Binary cross entropy loss for training and validation. The learned model was trained for blindly testing on chromosome 1. The validation data were extracted from chromosome 2. The training data were generated from the rest chromosome from 3 to X.

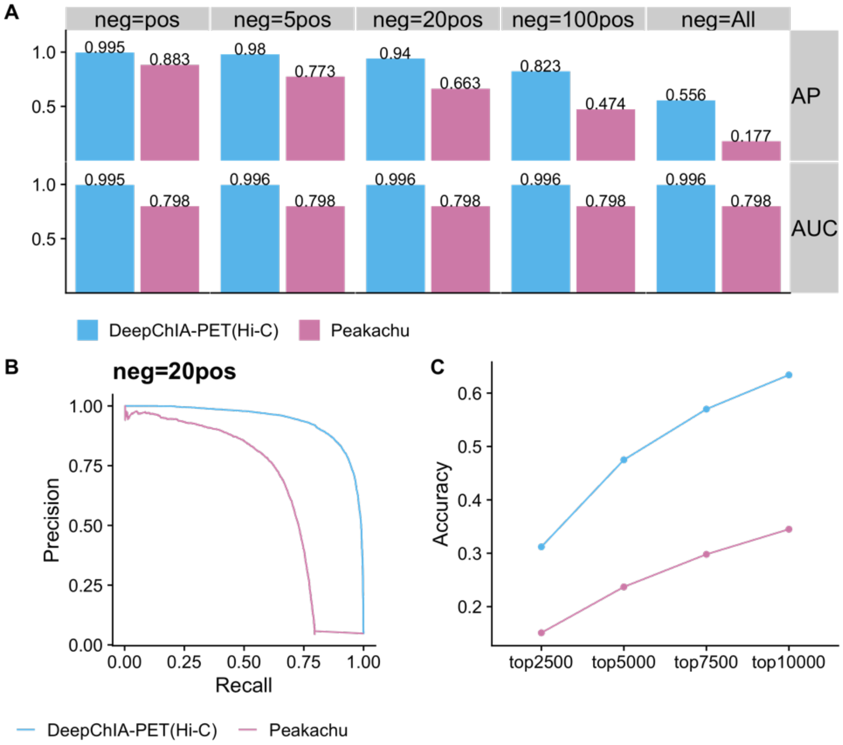


**Figure S4.** DeepChIA-PET(Hi-C) outperforms Peakachu for blindly testing CTCF ChIA-PET on chromosome 2 in GM12878. (A) The AP and AUC results for different number of negative pixels used for evaluating. (B) Precision-recall curve for the number of negative pixels equal to 20 times the number of positive pixels. (C) The topN accuracy where N equals four different values.

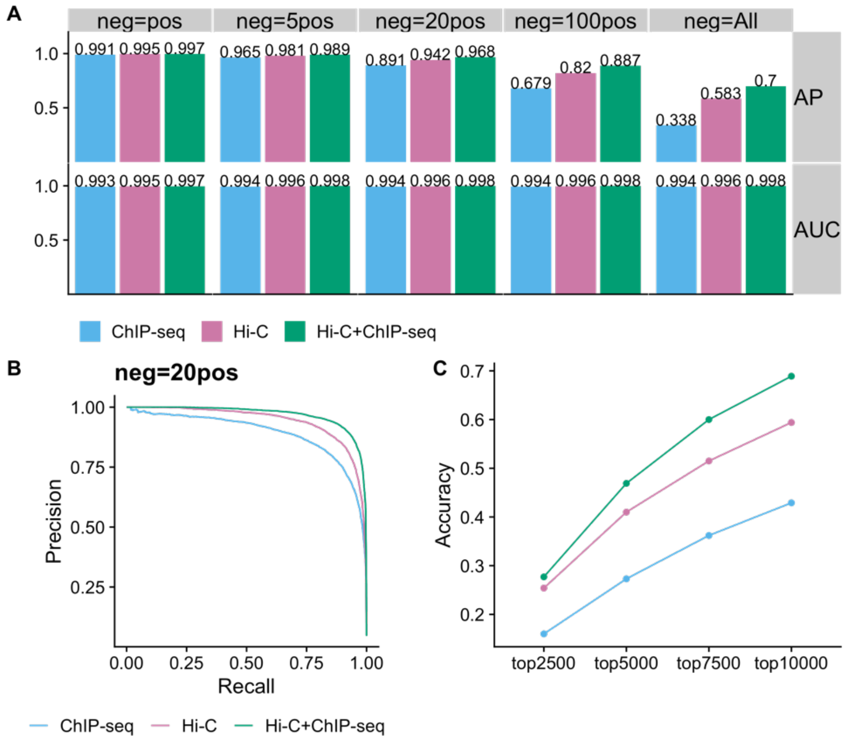


**Figure S5.** Adding ChIP-seq data as input improves the classification performance. The evaluation for DeepChIA-PET with only Hi-C, only ChIP-seq, and both as input is conducted on chromosome 1 for CTCF ChIA-PET in GM12878. (A) The AP and AUC results for different number of negative pixels used for evaluating. (B) Precision-recall curve for the number of negative pixels equal to 20 times number of positive pixels. (C) The topN accuracy where N equals four different values.

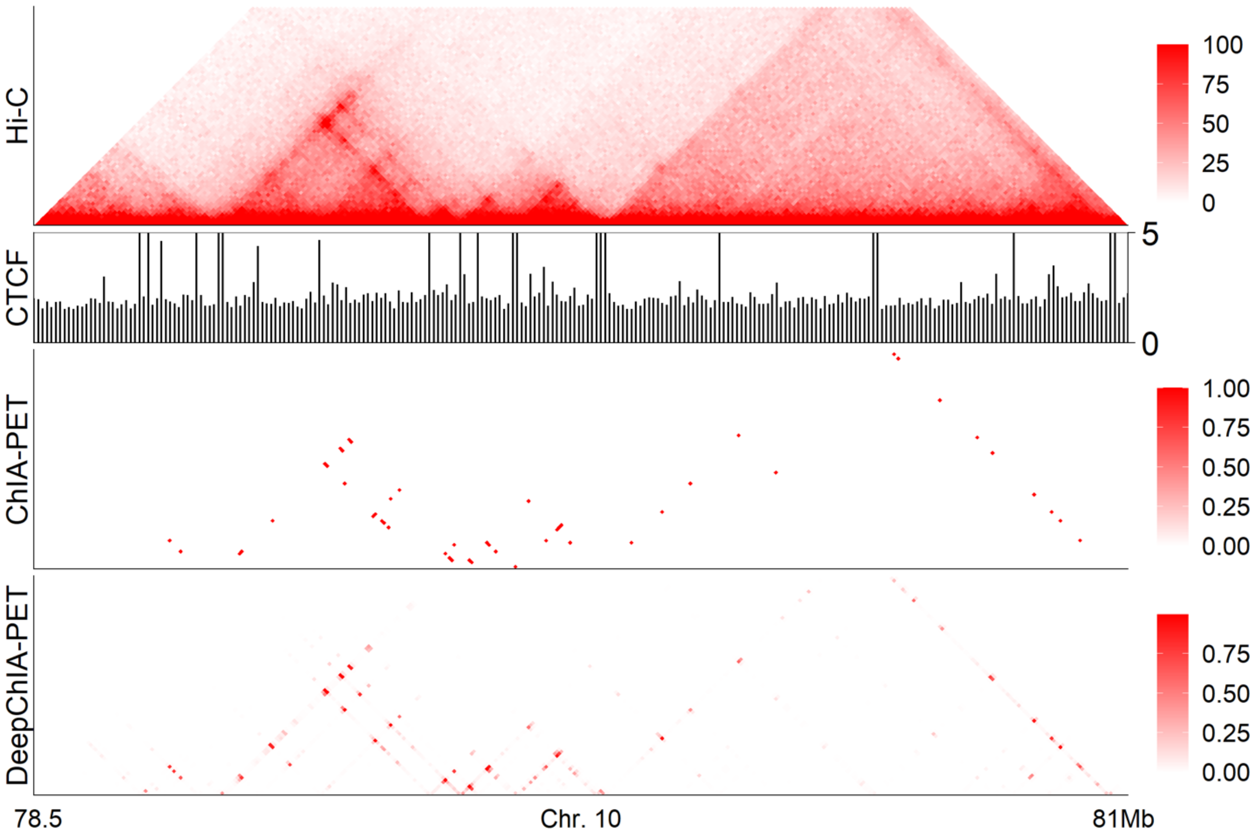


**Figure S6.** A specific example of our predictions for CTCF ChIA-PET interactions on chromosome 10 in GM12878. From top to bottom: KR-normalized Hi-C, CTCF, ground-truth ChIA-PET, and our predicted ChIA-PET.

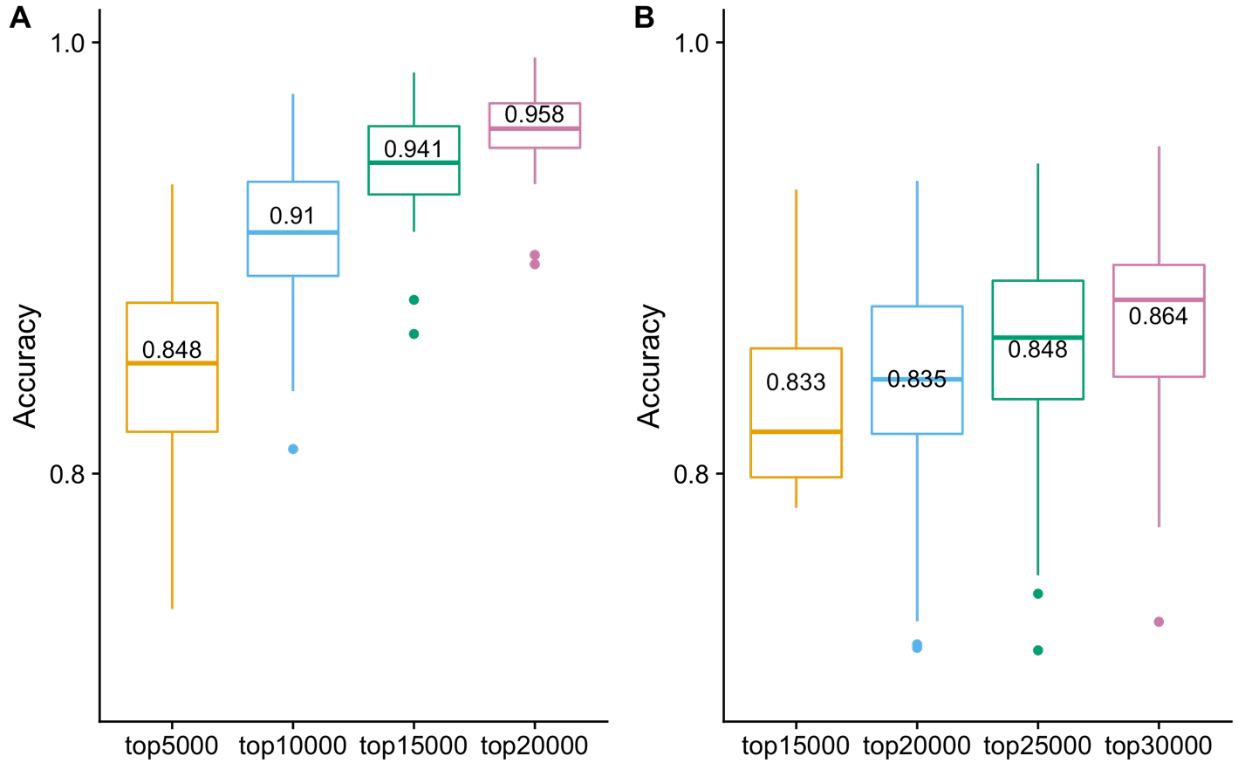


**Figure S7.** The topN accuracy scores of DeepChIA-PET for predicting RAD21 ChIA-PET of GM12878 (A) and K562 (B). Both (A) and (B) were generated on all testing chromosomes (mean values were added above each boxplot). Note: since the Hi-C matrix for chromosome 9 in K562 is empty, the evaluation results shown in (B) do not include the accuracy score for chromosome 9.
